# Dynamics and emergence of metachronal waves in the ciliary band of a metazoan larva

**DOI:** 10.1101/2025.02.10.637311

**Authors:** Rebecca N. Poon, Gáspár Jékely, Kirsty Y. Wan

## Abstract

Both natural and synthetic ciliary arrays exhibit diverse coordination patterns. Proper coordination of cilia is essential for the normal physiology of many organisms, from single cells to humans. Yet despite decades of research the mechanisms of cilia coordination remains disputed, particularly the question of how coordinated waves of activity known as metachronal waves arise in different ciliated systems. In many aquatic larvae that rely on ciliary motility to swim, the cilia are often arranged ornately in arrays or bands, within which robust metachronal waves propagate. Here, to resolve the origins of ciliary metachronism, we target the equatorial ciliary band of the marine annelid, *Platynereis dumerelii*, using whole-body high-speed imaging, and physical and biological manipulations. The results reveal an unprecedented wave structure featuring strong coupling within individual multiciliated cells that breaks down across cell boundaries, and absence of global coupling across the organism. Using laser ablation to create gaps in the ciliary band, we quantify the resulting disruption to wave propagation, revealing the extent of interciliary phase-locking and implicating steric interactions as an important contributor to coordination. The larvae also exhibit spontaneous whole-body ciliary arrests which allow us to study wave emergence and re-establishment for the first time, revealing a novel role of the animal’s nervous system in the dynamic coupling of cilia.

---

Cilia are hairlike organelles that are highly conserved throughout eukaryotes, in species ranging from humans to single-celled organisms [1, 2]. The primary function of motile cilia is to generate fluid flow, either for self-propulsion in freely moving swimmers, or to generate functional currents in the environment surrounding a sessile ciliated surface. Multiciliated cells and organisms can have as many as tens of thousands of cilia arranged in various geometries or topologies, which display diverse forms of coordination of their beat [3, 4]. Cilia beating can self-organise into different patterns including into rows of synchronously beating cilia with a constant offset between the beat phase of successive rows, forming *metachronal waves* [5, 6]. Wave ‘crests’ – where the cilia are at the start of their power stroke – propagate perpendicularly to the in-phase rows as the cilia beat. For all ciliated organisms, coordination of cilia is essential for the proper functionality of such arrays [7, 8]. These functions include self-propulsion in algae or marine larvae [9, 10], nutrient transport or fluid mixing in corals [11, 12], and mucociliary clearance where reduced coordination is associated with some forms of human primary ciliary dyskinesia [13, 14]. Ciliary metachrony has also been shown to enhance transport efficiency in magnetically actuated ciliary arrays [15, 16, 17], consistent with model predictions [18, 19, 20]. However, the mechanisms by which cilia achieve distinct forms of coordination in different systems are still not well understood [3].

Studies in several ciliated species have advanced our understanding of ciliary coordination and metachronal wave formation. Pairs of somatic cells isolated from the colonial alga *Volvox carteri* and held on separate micropipettes could synchronize their beating movements at distances below one cilium length, while the degree of synchronization decayed inversely with separation distance, consistent with hydrodynamic interactions [21]. In intact *Volvox* spheroids, ciliary metachronal waves are assumed to have a hydrodynamic origin [22]. In other systems however, including the biciliated protist *Chlamydomonas reinhardtii*, basal coupling dominates over hydrodynamic interactions [23, 24]. This type of coupling is attributed to calcium-sensitive filamentous connections in the basal apparatus [23, 25], with possibly homologous structures in other ciliates where they contribute to structural integrity, resist flow [26], or mediate gait coordination [27, 28]. Simplified mathematical and computational models have helped to interpret experiments and resolve the striking system-dependence of ciliary coordination, accounting for either hydrodynamic interactions [18, 29, 30, 31, 32], or basal coupling [33, 34], or both [35]. Coupled colloidal oscillators designed to mimic ciliary beating have also provided key insights into the relative importance of orbit shape and compliance [36]. Hydrodynamically coupled cilia do indeed exhibit metachronal waves in certain parameter regimes [18, 37, 38], but the case of high-density multiciliated arrays, more commonly found in metazoa, remains largely unexplored either from an experimental or theoretical perspective.

Here, in search of a suitable animal model to study ciliary coordination, we turn to the marine annelid worm *Platynereis dumerilii*, a powerful emerging model for neurobiology and systems neuroscience [40, 41]. The worms mature through a series of microscopic, planktonic larval stages before reaching their centimeter sized adult form (Fig. 1A). The trochophore larval stage of this organism occurs from 24-48 hours post fertilization (hpf), and has a prominent ciliary band called the prototroch. Arrangement of the cilia into dense stripes or bands is a defining morphological characteristic of many annelid species [5]. Trochophore larvae are planktonic and swim in helical paths (Fig. 1B). The larvae are spherical, with a diameter of ∼150μm, and the circular band of cilia (the prototroch) sits just above the larval equator [42] (Fig. 1C). Swimming is mediated by the concerted action of the many locomotor cilia and the larva’s neuronal circuitry, which takes input from sensory organs, such as eyespots. Synaptic input from the photosensitive eyespots to the ciliary band mediates larval phototaxis [43, 44]. Giant ciliomotor neurons that innervate the ciliary band cells drive spontaneous whole-body ciliary arrests, which regulate the position of the larva in the water column [45]. More generally, ciliary beating in marine larvae underlies many complex behaviours such as phototaxis, feeding, navigation towards suitable settlement sites, or escape from predators [46, 43, 47, 48].

**Figure 1:**
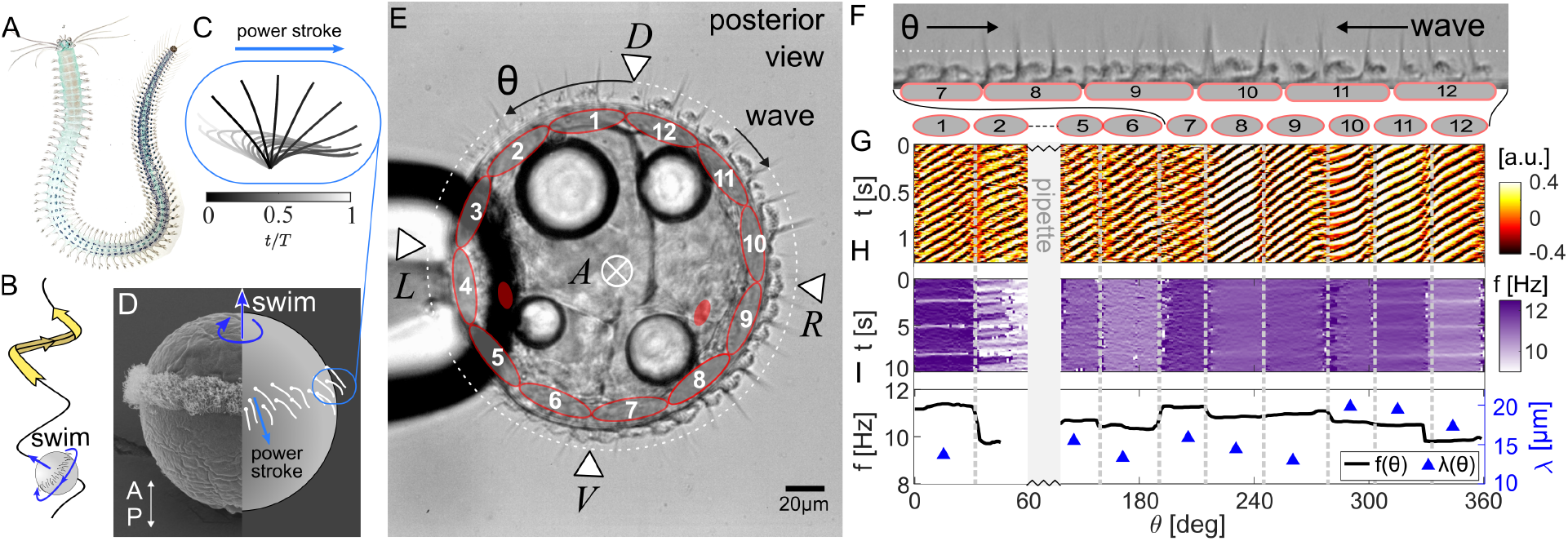
Spatial patterning in the dexioplectic ciliary metachronal wave of *Platynereis* larvae. (A) Image of an adult worm [39]. (B) The 24-hpf trochophore larva swims in a left handed helical path using cilia, which beat with a characteristic pattern (C). (D) Scanning electron microscopy (SEM) image of a 24-hpf larva spliced with a schematic showing axis orientation and swimming direction. (E) A single larva held by a micropipette with A-P, D-V axes labelled. The ciliary band comprises 12 large, distinct multiciliated ‘cells’ (see text), distributed regularly around the prototroch band. Two eyespots are marked in red. The angle *θ* around the band is defined with *θ* = 0 at the dorsal point. (F) An ‘unwrapped’ section of the ciliary band. (G) Kymograph of the image intensity around the ciliary band (dotted white line in (E,F)) as a function of *θ* and time *t*. Wave crests/troughs are visible as diagonal lines, travelling in the −*θ* direction. (H) Kymograph of the ciliary beat frequency as a function of *θ* and *t*, showing that the frequency is constant within each ‘cell’ but discontinuous over the cell boundaries (marked by grey dashed lines). (I) Time-averaged frequency and wavelength for this dataset.

This unique system gives rise to rich biophysics, allowing us to analyse in unprecedented detail the nature of ciliary beat coordination and the spatiotemporal dynamics of metachronal wave propagation and even emergence in a living specimen. The system exhibits a simple quasi-one-dimensional configuration with periodic boundary conditions, in contrast to the highly patchy or heterogeneous fields of cilia reported in other multicellular systems [49, 32]. It is known that the prototroch ciliary band exhibits metachronal waves [42, 45], but wave characteristics have not been quantified in detail. Here, we use micromanipulation to immobilise single larvae to visualise and characterise wave dynamics across the organism’s ciliary band under high-speed imaging. We discover a surprising *cell-scale* spatial patterning in the ciliary band metachronal wave. Using controlled laser ablation (for cilia removal) and pharmacological treatment (to modulate ciliary frequency) we perturb the system’s native ciliary coordination state, thereby decoupling the physical and biological factors that conspire to produce the final desired coordination outcome. Our results add substantial richness and novelty to the study of cilia coordination, emphasising the delicate interplay between strong local coupling within multiciliated cells mediated by physical interactions, versus global, neuronal coordination of the entire ciliary band.

## Results

### Ciliary band metachronal waves are spatially patterned

We focus exclusively on the early/mid-trochophore stage, 24–30 hours post fertilisation (hpf). Here, the prototroch ciliary band is composed of 23 multiciliated cells, 22 of which are arranged in stacked pairs (in the anterior-posterior direction), and one of which is unpaired [44]. The 11 pairs and one unpaired cell are distributed circumferentially around the ciliary band in 12 locations, numbered 1–12 starting from the dorsal point of the larva. Locations 1–11 each have one pair of multiciliated cells, with the unpaired cell at location 12. ^1^

Individual cilia are about ∼20μm long and 0.2μm in diameter. Due to their small size, ciliated microswimmers operate in the low Reynolds number fluid regime, where viscous forces dominate over inertia [50, 51]. The characteristic beat pattern comprises a power stroke, where the cilium is straight and fluid is pushed in the stroke direction, followed by a highly curved recovery stroke which returns the cilium to its starting position (Fig. 1D).

To visualise the ciliary band, individual larvae were aspirated onto micropipettes by gentle suction (see Methods). The larvae were repositioned with the full ciliary band focused in the imaging plane, and imaged using a high-speed camera at 100-500 fps (SI Video 1). Robust metachronal waves were observed to propagate circumferentially. We parameterise the distance around the band with angle *θ*, defined to be positive in the same direction as the cell numbering, with the origin *θ* = 0 at the dorsal point (Fig. 1E). We classify metachronal waves according to their propagation direction with respect to the power stroke [6, 5]: symplectic (if in the same direction), antiplectic (in the opposite direction), or diaplectic (if travelling perpendicularly). Diaplectic waves further differentiate into dexioplectic or laeoplectic depending on whether they travel 90^°^ anticlockwise or clockwise from the power stroke. Here we deduce that the ciliary band waves are *dexioplectic*, by plotting 2D kymographs of image intensity (function of position *θ*, and time *t*), which is a proxy for the ciliary beat phase *ϕ*(*θ, t*) [52]. Wave crests/troughs, as seen in the ‘unwrapped’ image of the ciliary band, appear as diagonal lines travelling in the −*θ* direction (Fig. 1F,G). The wave direction was dexioplectic in 100% of cases examined (n > 40 individuals). This directionality is consistent with observations in *Nereis pelagica*, a closely related species [6].

The kymographs show clear discontinuities spaced at ≈ 30^°^ intervals, which by inspection of the videos we confirmed were located at the natural boundaries between the 12 cell pairs (hereafter each referred to as a ‘cell’, for simplicity). Cell boundaries are subsequently located using discontinuities in the kymographs (see Fig. SI 1), and also visible in standard deviation projections of the ciliary band (see Fig. 4A). We calculated the spatially-resolved cilia beat frequency as a function of both space (position around the band) and time using a wavelet transform [53], plotted as a 2D map in Fig. 1F. The frequency profile is fairly constant over time (see SI for a longer dataset) but remains discontinuous in space. (Small frequency dips, seen as periodic horizontal white lines in the 2D frequency map, likely correspond to secondary neuronal activity, which we do not discuss further here.) Further, by time averaging the wavelet transforms to locate the peak frequency over 10 seconds, we see that the beat frequency is constant *within* each multiciliated cell, but discontinuous across neighbouring cells (Fig. 1G). Across the population (*n* = 21 larvae), the average beat frequency is 10 ± 2 Hz. The range of frequencies displayed within an individual larva is 1.6 Hz on average and can be up to 4 Hz. We also computed the metachronal wavelength (see Fig. SI 2), which exhibits the same piecewise continuous signature as the frequency. This robust spatial patterning was observed in all measured larvae. There was no clear correlation between a cell’s beat frequency and its anatomical location (see Fig. SI 3).

In summary, the *Platynereis* ciliary band propagates a robust dexioplectic metachronal wave. Yet, despite the illusion of a single globally continuous wave, there is in fact strong spatial patterning delineated by the physical boundary between neighbouring multiciliated cells. Within each cell, all the cilia adopt the same frequency, and a unified metachronal wavelength, but for cilia in different cells, this relationship breaks down. In other words, the cilia within the larva’s ciliary band perform a series of separate, *local* metachronal waves within each multiciliated cell.

### Cilia are phase locked within but not between multiciliated cells

So far, we have found frequency locking of cilia within, but not between multiciliated cells. Next we test for the stronger condition of phase locking. For this we construct a continuous phase *ϕ*(*θ, t*) from the image intensity timeseries by computing the Hilbert transform [54, 55]. This generates the imaginary analytic signal of a real time series, producing an instantaneous phase given by the polar angle. This phase grows from 0 to 2*π* over a full beat cycle (Fig. 2A), and can be unwrapped to remove jump discontinuities. If two oscillators with individual phases *ϕ*_1_(*t*) and *ϕ*_2_(*t*) are phase locked, their difference Δ*ϕ*(*t*) = *ϕ*_1_(*t*) − *ϕ*_2_(*t*) will be constant over time, otherwise, Δ*ϕ* will drift.

**Figure 2:**
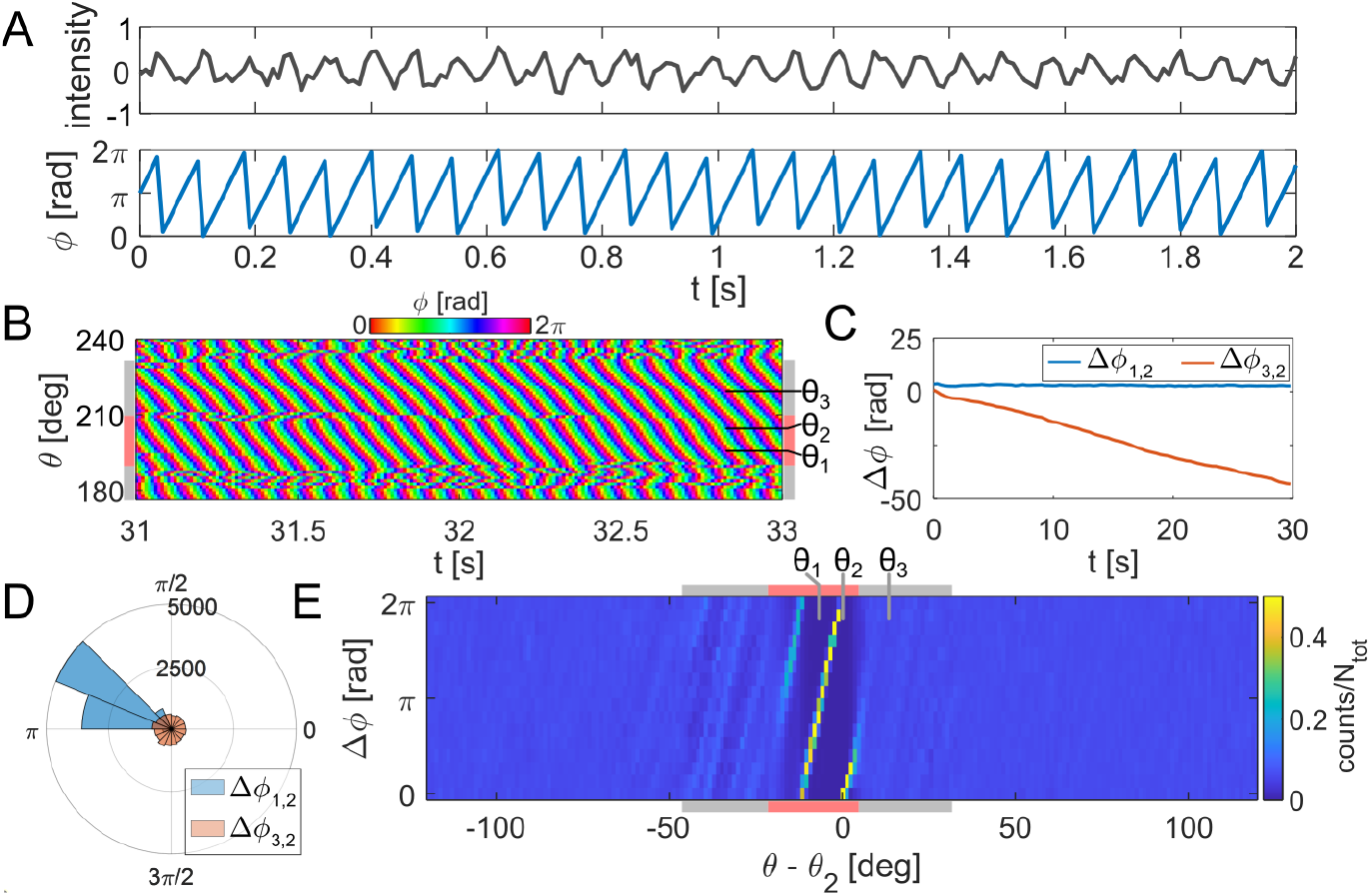
Ciliary beating is phase-locked within the same cell but not between cells. (A) Conversion of the intensity signal into a Hilbert phase *ϕ*, which increases linearly from 0 to 2*π* over one beat period. (B) The phase kymograph (similar to the intensity kymograph) across a region containing two cells in the ciliary band. Three distinguished locations are marked: *θ*_1_, *θ*_2_ in the same cell, *θ*_3_ in the adjacent cell. (C) Unwrapped phase difference Δ*ϕ*_*i*2_ = *ϕ*(*θ*_*i*_) − *ϕ*(*θ*_2_) for *i* = 1, 3, over 30 s. (D) Radial histograms of phase difference over the entire recording (100 s) showing that the cilia at *θ*_1_ and *θ*_2_ are phase locked, but *θ*_3_ and *θ*_2_ are not. (E) Plotting Δ*ϕ*_*i*2_ across the ciliary band shows strong phase locking only at locations within the same cell as *θ*_2_, not between cells.

For a sample larva, we plot *ϕ*(*θ, t*) for *θ* ∈ [180^°^, 240^°^], a region spanning approximately two cells (Fig. 2B). As expected, the phase is discontinuous across cell boundaries. At specific locations around the ciliary band, denoted by *θ*_1,2,3_ on the plot, we compute Δ*ϕ*(*t*) to investigate the phase relationship between pairs of cilia within the same cell (Δ*ϕ*(*t*)_1,2_ = *ϕ*(*θ*_1_, *t*) − *ϕ*(*θ*_2_, *t*)), or different, neighbouring cells (Δ*ϕ*(*t*)_3,2_ = *ϕ*(*θ*_3_, *t*) − *ϕ*(*θ*_2_, *t*)). We see that Δ*ϕ*(*t*)_1,2_ is constant over time, indicating cilia within the same cell are strongly phase locked (Fig. 2C). Meanwhile Δ*ϕ*(*t*)_3,2_ drifts steadily over time, even though in this instance the frequencies of two neighbouring cells are within only 2% of each other (at 12.8 Hz and 12.6 Hz respectively). This is also clearly seen by plotting the circular histogram of (wrapped) Δ*ϕ* over the 100 s time series; while Δ*ϕ*_1,2_ is peaked at a constant, but non-zero value, Δ*ϕ*_3,2_ is distributed uniformly (Fig. 2D).

We calculated Δ*ϕ* = *ϕ*(*θ, t*) − *ϕ*(*θ*_2_, *t*) at every location *θ* around the ciliary band with respect to the same reference location *θ* = *θ*_2_. This plotted as a 2D heat map (Fig. 2E). Where Δ*ϕ* is bounded (phase locked), the histogram shows a strong peak, coinciding with the interior of the reference cell. Wave propagation is detected as sharp diagonal lines. Outside of this region, Δ*ϕ* is spread over the full [0, 2*π*] interval, with traces of weak coupling with the nearest neighbours.

These results reveal that the cilia within each multiciliated cell not only have the same beat frequency but are also strongly phase locked, yet this coupling does not extend to cilia not within the same cell. Even when cilia from adjacent cells have a very small frequency difference, they still fail to phase lock across the boundary.

### Spatial gaps interrupt metachronal wave propagation

To understand why cilia do not phase lock across cell boundary, we must consider both the intrinsic beat frequency difference across distinct multiciliated cells, as well as the intercellular distance between cilia downstream of the previous cell and those upstream of the next. The coupling strength required for entrainment between the two groups of cilia will not be sufficient when the frequency difference is too high, and/or their separation is too large (especially since hydrodynamic interactions decay rapidly with distance [21]). Comparing the spacing between cilia within cells and across a cell boundary, we found that within each of the 12 cells, the cilia are spaced at an average distance of (0.57 ± 0.15)μm (a density of 3 ciliaμm^−2^), whereas the gap between two neighbouring cells is much larger, at (2.7 ± 0.8)μm (*n* = 4 larvae).

To test if lack of phase locking can indeed result from large gaps between cilia, we artificially introduced gaps of different sizes by controlled laser ablation *within* the naturally continuous ciliary field of a single cell (Fig. 3A). Fig. 3B,C shows a ciliary band before and after a gap has been introduced in the middle of a cell (SI Video 2). We confirmed by confocal immunofluorescence that cilia had indeed been removed from ablated regions (see Fig. SI 4). The procedure can even be performed multiple times on the same larva, as shown in Fig. 3(D–F). The ablated regions are clearly visible on intensity kymographs as zones with no periodic intensity signal, while ciliary metachronal waves persist on either side of the gap. As before, we compute the spatially-resolved beat frequency ⟨ *f* (*θ*) ⟩ (time averaged over 20 s). Comparing ⟨ *f* (*θ*) ⟩ profiles before and after the ablation, we find that ablation introduces a shift in the beat frequency of each multiciliated cell, but the beat frequency remains uniform within un-ablated cells, meaning that the cilia within each un-ablated cell remain locked to a common, but new, frequency. Within the ablated cell itself (e.g. Fig. 3D for a single ablation), a jump discontinuity develops across the gap so that the upstream and downstream groups of cilia are no longer locked.

**Figure 3:**
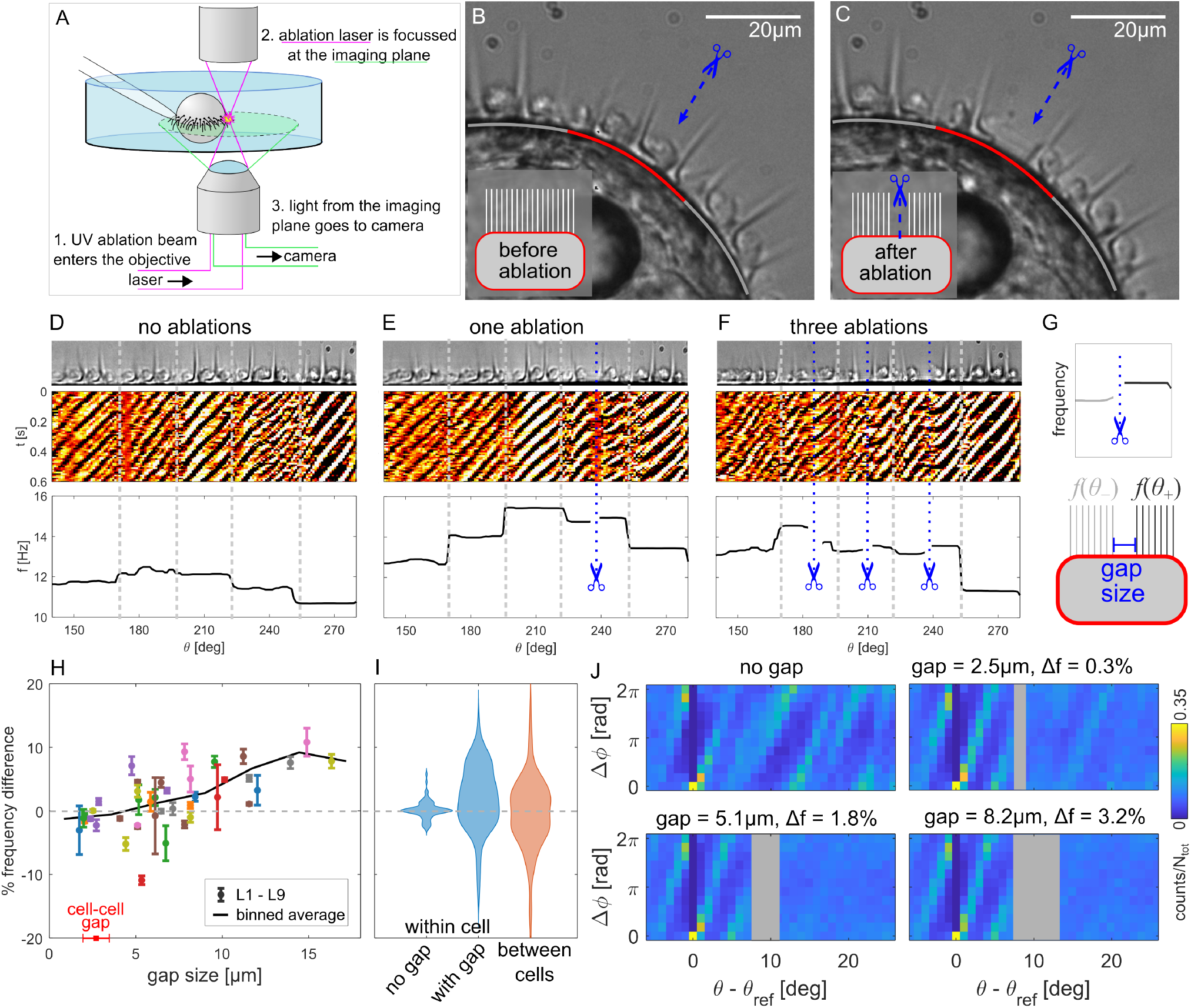
Metachronal waves do not transmit across small ablation gaps. (A) Schematic of laser ablation apparatus. (B,C) Still video frames showing the appearance of the metachronal wave before and after laser ablation. Grey-red-grey lines delineate the cell being ablated, and adjacent cells on either side. (D-F) Effect of successive laser ablations across 5 cells of a ciliary band. Top row: unwrapped view of the cilia; middle: corresponding intensity kymographs; bottom: spatially resolved frequency *f* (*θ*) profiles. Small gaps are ablated at the locations indicated. Cell boundaries are marked with dashed grey lines. (G) The two groups of cilia on either side of an ablation gap become decoupled, characterised by an upstream *f* (*θ*_+_) which differs from the downstream *f* (*θ*_−_) frequency. (H) The directional frequency difference Δ*f*_%_ across each ablation gap measured for 9 larvae (with 1 or more gaps per larva), also showing the approximate natural separation between adjacent ciliated cells. Each point shows the average and standard deviation for that gap over all available videos. (I) A violin plot of Δ*f*_%_ distributions within a cell, on either side of the ablation gap location before and after ablation; and between adjacent cells in the control data. (J) Set of histograms measuring Δ*ϕ* for a single cell before, and after three successive ablations of the *same* region (gap sizes of 2.5μm, 5.1μm and 8.2μm).

Next we investigate whether the cross-gap frequency difference is affected by ablation gap size. Each ablation is inherently asymmetrical since symmetry is broken by the wave which always travels in the −*θ* direction (Fig. 3G). We therefore define the cross-gap frequency difference by Δ*f* = *f* (*θ*_+_) − *f* (*θ*_−_), and its normalised version Δ*f*_%_ = 100 × Δ*f/f* (*θ*_+_). We plot Δ*f*_%_ against gap size (Fig. 3H), for data from n = 9 larvae and a total of 41 ablations, all for gaps below one cell-width (≈ 40μm). Both the average frequency and Δ*f* vary over the course of the experiment for each ablation gap, so for each gap size we plot the average value of Δ*f*_%_ over the entire experiment, with an error bar showing the standard deviation. The overall trend is that Δ*f*_%_ increases with increasing gap size. For the smallest gaps both positive and negative values of Δ*f*_%_ are possible, while large gaps (>10μm) always produce a higher upstream frequency after ablation. Fig. 3I summarises the distribution of Δ*f*_%_ measured within cells both before and after ablation, and between neighbouring cells in unablated (control) larvae. The intracellular frequency differences show a bias towards positive values (64% of all values are positive), whereas the intercellular values are symmetrically distributed around 0 (50% of all values are positive). The range of values is comparable.

We plot Δ*ϕ* histograms to evaluate the extent of phase locking within ablated cells, where 0.3% < Δ*f* < 3.2% (Fig. 3J)). These show that the cilia generally remain phase locked within the patch on one side of the gap, but are not strongly coordinated across the gap, even when the frequency difference across the gap is very small (comparable to the frequency noise of 1.5% within the intact patch).

Finally, to establish whether global cilia-generated flows could influence coordination within the band, we progressively ablated large regions of the ciliary band until only ≈ one wavelength of the wave remained (SI Video 3). Figure 4 shows kymographs, frequencies and wavelengths of the same region, featuring one cell, then only a decreasing patch of cilia, during successive ablations. The wave continues almost completely unchanged in the small region even as more and more cilia in the band have been removed.

**Figure 4:**
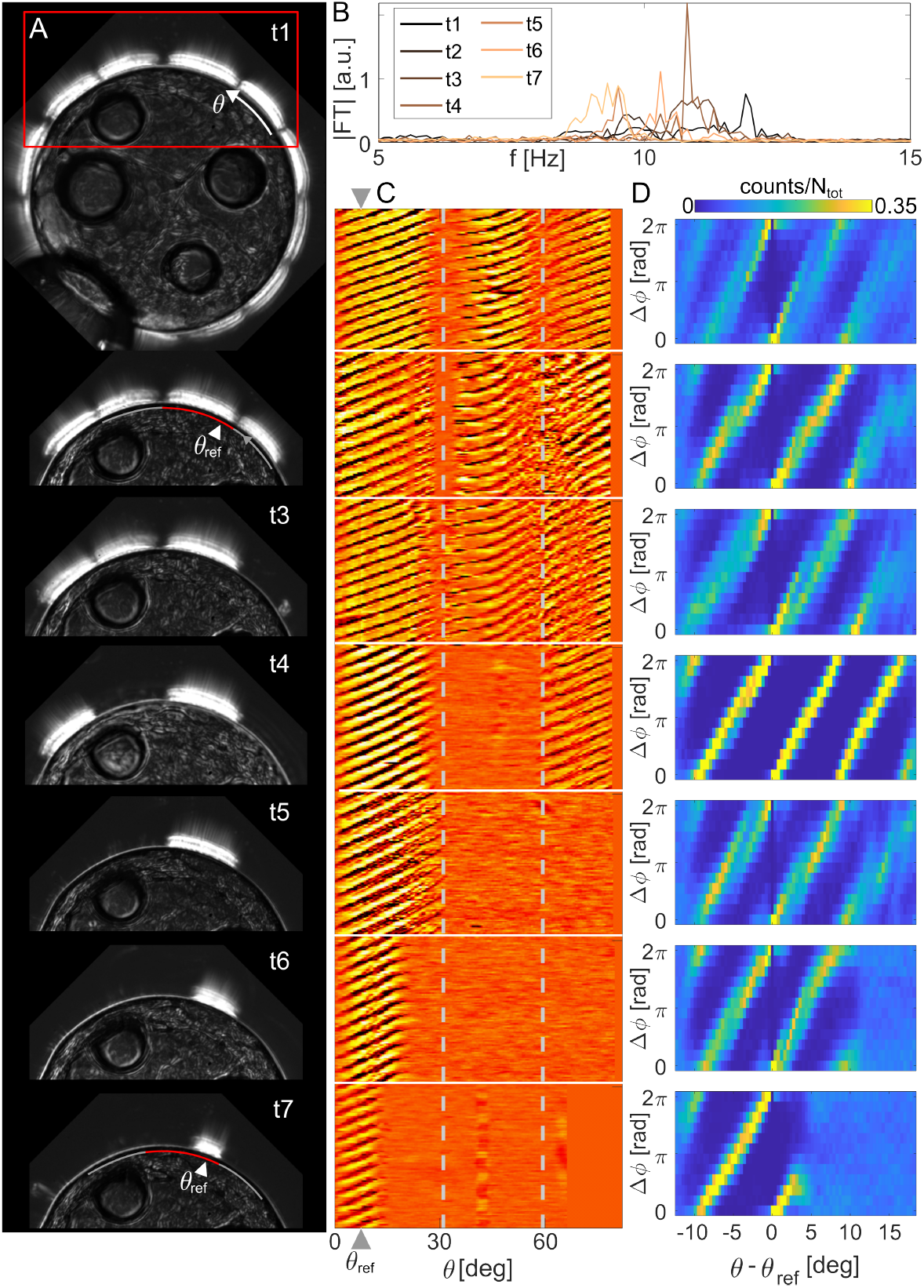
Metachronal waves are locally robust to large-scale ablation of the ciliary band. (A) Standard deviation projections of the relevant section of the ciliary band (remaining cilia are visible as bright patches), over 7 time points representing successive ablation events. (B) Fourier transform of the intensity signal, averaged over the remaining cilia for the cell of interest. (C) Intensity kymographs over the region of interest, measured after successive ablations. (D) Histograms of Δ*ϕ* for only the highlighted cell, showing minimal variation in coordination, frequency and wavelength within the remaining patch over the entire course of the experiment, showing that *local* wave propagation is independent of other cells or global ciliary flow.

Thus, the ciliary band metachronal wave is interrupted by even small spatial gaps within the same cell. Yet, patches of cilia left intact by ablation still remain frequency and phase-locked to the ‘self’ metachronal wave, completely independently of cilia elsewhere in the other cells.

### Metachronal wave arrest and re-emergence

In addition to the metachronal wave itself, *Platynereis* larvae exhibits spontaneous ciliary arrests known as closures [45]. This natural behaviour is used by the larvae to regulate their vertical position in the water column [45]. Here, we leverage this physiological response to investigate how ciliary metachronal coordination is *established* in a living system for the first time.

SI Video 4 shows an example of a closure in a pipette-held larva with the entire ciliary band visible. To identify the closure events, we first condense the 2D kymograph information into a 1D time series, by defining a coordination number *χ*(*t*),

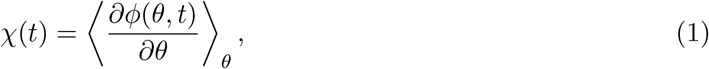

where *θ* is the angular distance along the array, and ⟨…⟩_*θ*_ denotes the average over all positions in the array, and *ϕ* is the instantaneous phase introduced above. When all of the cilia in the array are beating either randomly or synchronously, then *χ* = 0, whereas for a metachronously coordinated array *χ* will take a finite positive value. During periods of coordination, when *χ* is given in units of [rad/deg] (Fig. 5), the wavelength λ in ‘degrees around the band’ is given by λ = 2*π/χ*, or alternatively in units of length by 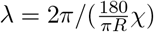, where *R* is the radius of the larva, *R* ≈ 75μm.

**Figure 5:**
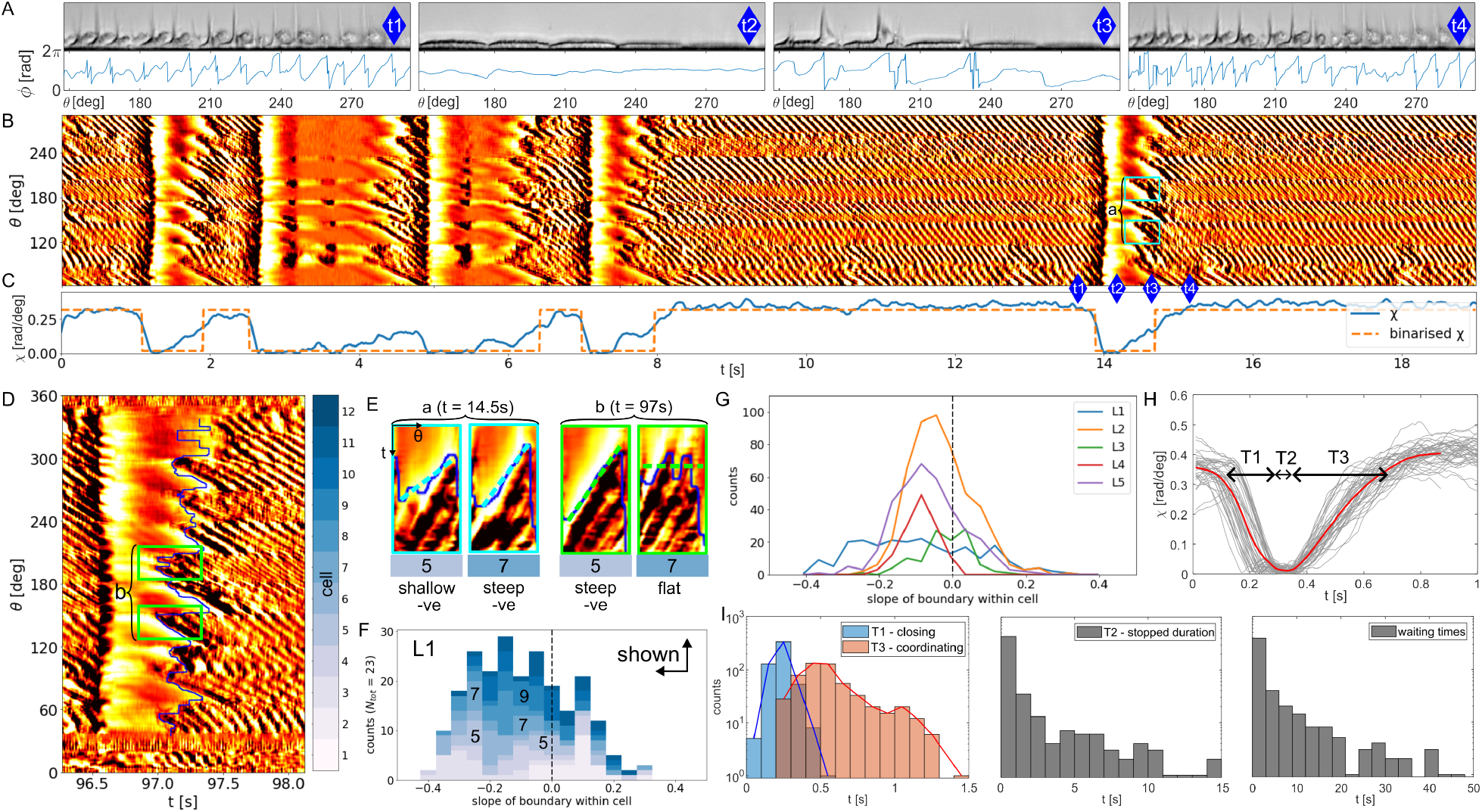
Emergence of metachronal waves after whole-body ciliary arrest. (A,B) Closures are a physiological response where the entire ciliary band stops beating and then resumes over time (SI Video 4). Snapshots shown at four distinct times during a typical closure, marked on the kymograph, with the corresponding unwrapped phase. The cilia beat normally at (t1), are fully arrested at (t2), resume beating at (t3), recover the final coordinated wave pattern at (t4). (C) The coordination index *χ*(*t*) is used to identify closures automatically. (D) Zooming in on the highly stereotyped dynamics during a single closure. We first locate the boundary between ‘closed’ and ‘beating’ cilia over the whole ciliary band. For each cell, we measure the slope of the recovery boundary (small panels on the right). (E) A flat slope means that the cilia restart nearly simultaneously across the whole cell, while a strongly negative slope indicates cilia start beating from the ‘upstream’, +*θ* side of the cell and progressively restart downstream. This behaviour is variable over time, and across cells (compare Cells 5, 7 ad 9 (F) Histogram of slopes for all 12 cells of the ciliary band, for the larva shown in (a), colour coded by cell number. (G) Distribution of boundary slopes for 5 different larvae. (H) Successive closures for the same larva are stereotyped, allowing us to define 3 timescales, *T* 1 for the time to arrest all cilia from a wave-state, *T* 2 the duration of the stopped phase, *T* 3 from the time the beating restarts to fully established wave. (I) Distributions of *T* 1 − 3 (n=9 larvae) and waiting times (duration between the end of one closure and the start of the next). For reference, the average ciliary beat period is 0.1 s.

Fig. 5A,B follows a series of spontaneous closures in a single larva. The intricate structure of metachronal wave death and re-emergence are revealed in the intensity kymographs. Closures, which can be detected automatically by thresholding the corresponding *χ*(*t*) timeseries (Fig. 5C), exhibit a high degree of stereotypy, but also some heterogeneous features. At onset of a closure, all cilia stop beating almost simultaneously, consistent with neuronal signalling. The cilia are usually arrested for a brief period before resuming their beating at slightly different times along the band, often with some spatiotemporal patterning already present. The recovery profile is more variable, the metachronal wave usually takes a few beats (5 ∼ 10) to reach steady state, and its exact progression can vary significantly across different larvae, from cell to cell, and even from closure to closure within the same larva.

Surprisingly, in a subset of the cells ciliary beating re-starts at the upstream end of each cell and propagates downstream. Zooming in on a single closure (Fig. 5D), we can quantify wave re-establishment at the cell-level, focusing on the slope of the ‘boundary’ between where the cilia are closed and when they are beating. A flat slope means that the cilia restart nearly simultaneously across the whole cell, while a strongly negative slope indicates cilia start beating from the ‘upstream’, +*θ* side of the cell and progressively restart downstream. To quantify the variability within the same individual over time, we plot a stacked histogram of the per-cell slope boundaries for a larva which shows a ‘mixed’ behaviour, differentiating between measurements from the 12 cells that constitute the ciliary band (Fig. 5F). Across all measured individuals (*n* = 5), there is a notable bias toward negative slopes (Fig. 5G).

We quantified the time taken to recover from the low closure value to the high value associated with a metachronal wave. Closures within a single individual are also stereotyped, allowing us to define three distinct timescales, *T* 1 for the time to arrest all cilia from a wave-state, *T* 2 for the duration of the stopped phase, and *T* 3 from the time the beating restart to fully established wave. Fig. 5H. We plot the resulting distribution in Fig. 5I. The ‘closing’ time *T* 1 and ‘coordinating’ time *T* 3 are narrowly distributed, peaking at 0.3 and 0.5 respectively, whereas the arrested timescale *T* 2 is much more variable. In some cases, the cilia can remain arrested for many seconds (SI figure). Finally, we also plot the waiting time between successive closures (here measured from the end of one closure event to the start of the next).

Our analysis of whole-body ciliary closures revealed that neuronally triggered ciliary arrest was nearly instantaneous and simultaneous across across the entire ciliary band, whereas the reestablishment of the metachronal wave was more variable and could take several beat cycles to complete across all cells. Importantly, the re-establishment of coordination displayed unidirectional spatiotemporal patterning, contrary to predictions of existing computational models (see Discussion).

### Spatial variation in serotonin response

Next, we explore the dependence of ciliary beating and metachronal wave properties on perturbations of biochemical origin, with exogenous addition of the neurotransmitter serotonin. In many organisms, serotonin increases the beat frequency of the cilia [45, 5, 56, 57]. Here, it acts on serotonin receptors expressed in the multiciliated cells via a G-protein cascade that leads to cAMP increase. We imaged the ciliary band activity of *Platynereis* larvae while adding different concentrations of extracellular serotonin. Briefly, successively higher concentrations of serotonin (from 0 to 50μM) were introduced to the medium in the vicinity of a pipette-held larva (see Methods).

A discontinuous frequency profile was again observed in all cases. Interestingly, increasing serotonin concentration appeared to shift the entire ciliary band frequency profile upwards to higher frequencies (see Fig. 6A,B for two examples of this). This shift seems to saturate at the highest concentrations, and not every cell displayed the same frequency response. For a subset of experiments, the medium was flushed out after the highest concentration, confirming that the response was reversible and the original frequencies could be regained. However, the ‘washed out’ frequency profile was slightly higher than the original one (see Fig. SI 5), implying that the intracellular concentration may still be higher than the resting value and will take longer to fully equalise.

**Figure 6:**
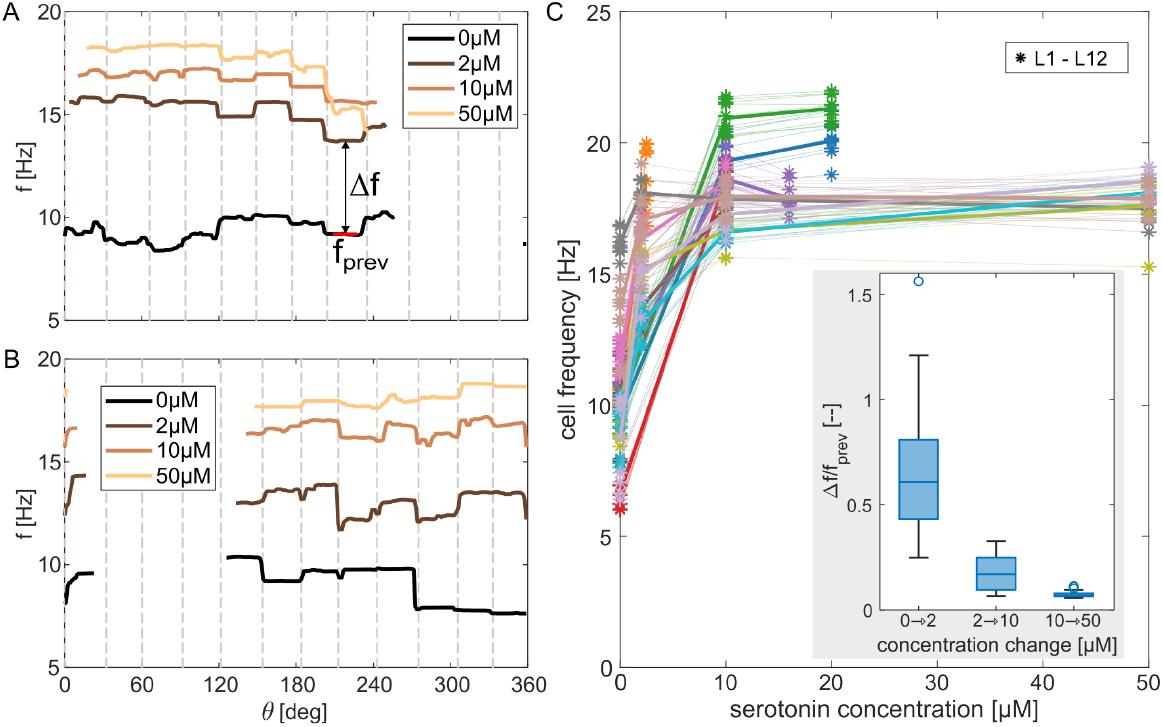
Response of the ciliary band metachronal wave to serotonin. (A) Measured *f* (*θ*) across the ciliary band for the same larva at different concentrations of exogenous serotonin, including the control. The profile changes the most between the initial value and the first concentration point, but thereafter retains its characteristic shape. (B) Example plotted for a second individual. (C) The per-cell beat frequency increases as a function of successive changes in serotonin concentration (for *n* = 6 larvae), but saturate at the highest concentrations. Inset: the ‘initial’ frequency jump is large compared to the subsequent frequency increases, as is the variation in the data.

We also quantify the per-cell beat frequency change as a function of serotonin concentration (Fig. 6C), for 12 individuals, excluding any ‘washed out’ data points. The response curve saturates at the highest concentrations. Although the resting frequencies range from 7−16 Hz, the frequencies all reach a maximum value of (18.6 ± 3.7) Hz, with the plateau beginning at around 10μM.

These results show that the overall frequency profile for each larva is highly robust. Across the population, the animals responded to increasing serotonin by increasing the ciliary beat frequency, until the point of saturation, with evidence of individual ciliated cells retaining their ‘identity’ throughout. This response is reversible, with cilia recovering the control dynamics after removal of the stimulus.

## Discussion

The capacity for multiple biological components to come together to produce emergent functions is one of the most fundamental features of active living systems. Here, we investigated how thousands of cilia couple across different spatial and temporal scales to produce coordinated waves of activity. This universal phenomenon is critical to the natural swimming behaviour of many marine zooplankton to control their ciliary beating and to regulate their position within the water column. Our results show that the locomotor cilia of *Platynereis* perform a robust metachronal wave that is interrupted by periods of spontaneous ciliary arrest known as closures. Surprisingly, we found that despite the illusion of a single continuous wave, the ciliary band metachronal wave is in fact a concatenation of locally metachronal waves each persisting only over the scale of a single cell. This unique system has generated fundamental insights not only into how cilia are coupled but also the re-establishment and control of global patterning across the entire ciliary band.

The robust spatial frequency profile across the ciliary band reveals strong local coupling and absence of global coupling. Cilia within each cell are tightly coupled to the same frequency and phase-locked, yet cilia across the small boundary between cells completely fail to synchronize. There was no phase-locking even over distances of ∼4μm (cilium length *L* ≈ 20μm), neither across natural gaps at cell boundaries, nor artificial gaps cut within cells by laser ablation – patches of cilia on either side of an ablation gap remaining strongly self-coupled. This strongly local nature of ciliary coordination is most strikingly demonstrated in ablation experiments where we removed all except a small patch of cilia on the entire band, which continued to propagate an almost unchanged wave pattern. Global flow therefore has no effect on the coordination. Ciliary phase-locking evidently does not require the entire cell to be intact, only that the cilia are spatially continuous. We deduce that short-range physical interactions between the cilia must be responsible for their strong local coupling. This interaction must also be spatially fast-decaying, given that metachronal waves fail to propagate even across very small ∼4μm gaps within the same cell, even with a negligible cross-gap frequency difference. Operating over such short distances are steric hindrance, hydrodynamic interactions and basal coupling by cytoskeletal fibres [8, 23]. Since our laser ablation experiments preserve internal connections between the cilia, this leaves steric or hydrodynamic interactions as the key contributions most affected by artificial gaps within cells.

In hydrodynamically interacting cilia arrays, diaplectic metachronal waves rarely emerge: instead antiplectic or symplectic metachronal waves are most commonly seen in simulation as well as synthetic and natural systems [21, 20, 58, 59, 60]. This is likely due to weaker hydrodynamic forces between cilia arranged in a diaplectic row (oriented with their principal beat planes perpendicular to the row), compared to when they are aligned with their power strokes in the same plane. For a 1D chains of cilia this interaction is estimated to be 100× weaker [61], ensuring a fast spatial decay. The ciliary band metachronal wave exhibits further features that are inconsistent with predictions in purely hydrodynamically-coupled ciliary arrays, notably the metachronal wavelength, and timescale for wave establishment. Here, the diaplectic wave has a short wavelength of ≈ *L/*2, compared to longer wavelengths ≳ 2*L* in previous model predictions. Even though the process is noisy, the timescale for local wave emergence as soon as the cilia start to beat (sometimes with immediate spatial patterning) is shorter than estimates from hydrodynamically coupled arrays (10^3^ beat cycles in some cases) [37, 22]. Therefore, while they contribute to local coupling, hydrodynamic interactions alone would appear to be insufficient in our case. Instead steric interactions, inevitable between densely packed cilia, must play a crucial role. This is neglected in most simulations that enforce a separation of at least 1*L* − 2*L* between cilia to avoid collisions, which also sets a lower bound on the metachronal wavelength of ≳ 2*L* [37, 62]. We additionally note that these minimum wavelengths found in simulated arrays correspond to approximately antiphase synchronisation of neighbouring cilia, which is qualitatively different to what is observed in our system. Indeed metachronal waves induced by local steric repulsion in arrays of filaments have been observed in hybrid molecuar dynamics simulations [63], as well in natural systems of free swimming nematodes at high density induce[64]. Thus, despite their ubiquity in many naturally-occurring systems [5, 10, 32, 65], diaplectic metachronism clearly shows very different features to anti- or symplectic waves that are not well captured by current formalisms.

Our study addresses the key open question of how the ciliary metachronal wave direction is selected in different systems [8, 6, 32]. Here, the waves were dexioplectic in 100% of the larvae we examined, this extraordinary directional robustness implicates existence of directional biochemical signalling, some underlying physical asymmetry in the ciliary array itself, or both. The first possibility seems unlikely: as the wave emerges during closures, local patches of beating cilia assumed a wave-like pattern in the same direction as the fully-established wave almost immediately (within one beat period < 100ms). Although ciliary beating restarts after a closure in response to a fall in the intracellular calcium concentration (as shown by calcium imaging in [45]), the timescale for calcium to diffuse across a cell (estimated using measured diffusion rates of Ca^2+^ ions in different environments [66]) is much longer than that associated with onset of spatial patterning, which is ≲ 0.5 s. This effectively rules out local gradients in intracellular calcium signalling *along* a given multiciliated cell. An essential clue lies in the larvae’s slow, chiral self-rotation around the AP axis as they swim [43]. The ciliary waveforms cannot be perfectly parallel to the AP axis, or must have a 3D component [67, 68, 69]. We estimate a tilt of 10°-15° of the ciliary power stroke in the −*θ* direction by direct measurement of flow fields around a pipette-held larva, making the wave slightly dexio-symplectic. This physical asymmetry will introduce directional bias in both the steric and hydrodynamic interactions between the closely packed cilia in the prototroch. This inherent tilt, alongside the gradient we measured in the intrinsic beat frequency across each cell, may be responsible for selecting the dexioplectic rather than laeoplectic wave [70, 71]. By accessing the intrinsic frequencies of the two groups of cilia on either side of an ablation gap, we found that for the largest gaps, where cilia are expected to be almost completely decoupled, beat frequencies were as much as 10% higher upstream of the ablated cell than downstream. Indeed, for a linear array of coupled Kuramoto oscillators, gradients in intrinsic frequencies of the oscillations promote metachronal wave propagation from the higher towards the lower frequency oscillators [72]. This putative relationship between 3D ciliary beating, intrinsic frequency gradients and wave propagation direction remains to be examined in other organisms.

In all multiciliated systems, achieving proper coordination is essential for their physiological function, whether this is coordinated swimming in ciliates, or mucociliary clearance in ciliated epithelia [73, 49, 74, 75]. Here in the ciliary band of invertebrate larvae, we found a remarkable metachronal wave construction comprising locally continuous waves within cells, with nervous signalling (for example through neuropeptide release) as the sole origin of global control over the entire band. Since the larvae rely on the ciliary band for motility and predator avoidance [46], this unique hierarchical design makes the wave extremely robust to accidental damage or loss of cilia. The *function* of the ciliary band is thus largely immune to local defects in wave propagation. This modular design effectively decouples the local scale at which self-organised interactions act (decentralised) from the large-scale neuronal or electrochemical control over the entire organism’s body (centralised). Similiar measures to safeguard biological robustness may already be present in early-diverging animals such as Placozoa [76], even in unicellular organisms with complex infraciliature [77] or ciliary bands in *Stentor* and *Didinium* [52, 74].

## Methods

### Animal husbandry

Platynereis dumerilii larvae were obtained by spawning one mature adults male and female derived from a culture at the University of Exeter. Cultures were maintained in temperature-controlled (22 +/- 2 C) rooms in plastic boxes (Vitlab 36491) containing 1.5 L steril artificial sea water (Tropic Marin Pro-Reef salt) at a salinity of 33 ppt. We provided a 16:8 hr light:dark photoperiod, with an artificial moon cycle (30 % light intensity overnight for 6 continuous nights within each 28-day cycle). Diet included live microalgae, live rotifers, and spinach, as described in (Hird et al). After spawning one male and one female epitoke, fertilised eggs were rinsed and embryos developed in artificial sea water in an 18 °C incubator under a 16:8 hr light:dark light cycle. Please refer to the detailed protocol in Hird et al [78].

The larvae were grown to an age of 24 hpf, and all data in this paper was taken using larvae between 24 and 34 hpf.

### Imaging and micromanipulation

For live imaging, larvae were held on a glass micropipette, mounted on a Scientifica PatchStar micromanipulator turret. The pipettes were pulled from TW-120 borosilicate glass (World Precision Instruments) using the ‘long taper’ program on a Sutter P-1000 pipette puller, then scored, broken back and fire polished (see Fig. SI 6 for details) to give an inner diameter of 20 − 30μm and wall thickness of 30 − 40μm, with rounded walls to minimise the damage to the larval tissue whilst being held. Larvae were placed in an imaging petri dish (Ibidiμ-Dish 35 mm, low, uncoated) in ≈800μl of seawater, then caught in an appropriate orientation by repeated release/capture cycles using the glass pipette. For measurement of the metachronal wave, the larvae were caught in the natural vertical swimming orientation, with the prototroch ciliary band in the imaging plane and the direction of swimming pointing upwards. The pipette and dish were manoeuvred until the larva was close to the centre of the droplet, in order to minimise boundary effects. Imaging was performed using a Leica DMi8 inverted microscope, at various magnifications, and videos were captured using a Phantom V1212 high-speed camera.

### Serotonin experiments

Videos were taken for 3 minutes at 100 fps, then for 10s at 500fps, then a small volume of stock serotonin solution was added to the imaging dish. The larvae were left for 5 minutes to allow uptake of the serotonin, then imaged again for 3 minutes at 100 fps and 10s at 500fps. This process was repeated to give multiple increasing concentration points.

### Image analysis and parameter estimation

Kymographs of the image intensity as a function of time were produced for a region along the full circle of the ciliary band [52]. Frequencies were calculated for each row of the kymograph (corresponding to the *I*(*t*) for one pixel along the band), using a wavelet transform to give time-resolved values of the frequency spectrum. The time-average of the frequency spectrum at each location was used to find the peak frequency values as a function of *θ*. The wavelet transforms were carried out using the MODA toolbox in Matlab [53].

### Laser ablation

Laser ablation was performed using a UGA-42 Caliburn pulsed 355nm UV laser from Rapp Opto-Electronic. The laser was set up on an Olympus IX83 inverted microscope, and videos were taken using a 01-Kinetix-M-C camera or an Excelitas pco.panda 4.2 camera. To perform the ablation, the larvae were held on a glass micropipette with the ciliary band oriented in the imaging plane. The laser was then programmed to fire continuously at 1000 Hz in a user-defined line somewhere within the ciliary band, and the focus of the laser was moved up and down to ensure that cilia within the entire height of the band were ablated. This process normally took around (20 ± 10) s. The laser power was attenuated by an ND2 filter (giving a 100× power reduction), and further set to fire using between 50% and 100% of the attenuated power, in order to ablate the cilia reliably without causing shock damage to the larva. The absence of cilia in the ablated gap was confirmed by visual inspection before further measurements were taken. For a single larva, a number of cells could be ablated in this fashion with minimal effect on the behaviour of the other cilia around the body. The gap sizes were measured from the video data. If possible, they were measured during a closure, where the gap is more clearly visible. We confirmed the effectiveness of this protocol via confocal imaging of larval cells whose cilia had been completely ablated. The larvae were stained for acetylated tubulin, and imaged in fluorescence to allow the cilia to be clearly viewed (see Fig. SI 5). Most of the cilia have been ablated completely, leaving only a very short region embedded in the cell, visible as well defined points in the image. Some cilia have been partially ablated so that short stumps of <5μm length remain.

## Acknowledgments

This work was funded by the European Research Council (ERC) under the European Union’s Horizon 2020 research and innovation programme grant 853560 EvoMotion (K.Y.W), grant 101020792 PROTOEYE (G.J.). This work was also funded by the Wellcome Trust (214337/Z/18/Z). For the purpose of open access, the authors have applied a Creative Commons Attribution (CC BY) licence to any Author Accepted Manuscript version arising.

We thank Luis Bezares Calderón for preparing and imaging the tubulin stained larvae following laser ablation, and for sharing his extensive *Platynereis* knowledge. We thank Adam Johnstone and the Marine Invertebrate Culture Unit at University of Exeter for providing *Platynereis* cultures.

## Footnotes

1 The numbering convention in the literature is still being standardised: here we follow the literature convention of the numbers increasing in the clockwise direction when viewed from the anterior (so, anti-clockwise when viewed from the posterior as in most of our videos), but number them so that the dorsal point sits between locations 12 and 1. This is different to the numbering given in [43] and is rotated by one location compared to the numbering in [45], which numbers the unpaired cell as 11.

## References

[1] Mitchell, D. R. The evolution of eukaryotic cilia and flagella as motile and sensory organelles. Eukaryotic membranes and cytoskeleton: Origins and evolution 130–140 (2007).

[2] Gibbons, I. Cilia and flagella of eukaryotes. The Journal of cell biology 91, 107s–124s (1981).

[3] Wan, K. Y. Coordination of eukaryotic cilia and flagella. Essays Biochem. 62, 829–838 (2018).

[4] Gilpin, W., Bull, M. S. & Prakash, M. The multiscale physics of cilia and flagella. Nature Reviews Physics 2, 74–88 (2020).

[5] Marinković, M., Berger, J. & Jékely, G. Neuronal coordination of motile cilia in locomotion and feeding. Phil. Trans. R. Soc. B 375 (2020).

[6] Knight-Jones, E. W. Relations between Metachronism and the Direction of Ciliary Beat in Metazoa. Journal of Microscopic Science s3-95, 503–521 (1954).

[7] Párducz, B. Ciliary Movement and Coordination in Ciliates. Int. Rev. Cytol. 21, 91–128 (1967).

[8] Wan, K. Y. & Poon, R. N. Mechanisms and functions of multiciliary coordination. Current Opinion in Cell Biology 86, 102286 (2024). URL 10.1016/j.ceb.2023.102286.

[9] Ishikawa, T. Fluid dynamics of squirmers and ciliated microorganisms. Annual Review of Fluid Mechanics 56, 119–145 (2024).

[10] Poon, R. N. et al. Ciliary propulsion and metachronal coordination in reef coral larvae. Physical Review Research 5 (2023).

[11] Shapiro, O. H. et al. Vortical ciliary flows actively enhance mass transport in reef corals. Proceedings of the National Academy of Sciences 111, 13391–13396 (2014).

[12] Pacherres, C. O., Ahmerkamp, S., Koren, K., Richter, C. & Holtappels, M. Ciliary flows in corals ventilate target areas of high photosynthetic oxygen production. Current Biology 32, 4150–4158 (2022).

[13] Lechtreck, K.-F., Delmotte, P., Robinson, M. L., Sanderson, M. J. & Witman, G. B. Mutations in hydin impair ciliary motility in mice. The Journal of cell biology 180, 633–643 (2008).

[14] Meeks, M. & Bush, A. Primary ciliary dyskinesia (pcd). Pediatric pulmonology 29, 307–316 (2000).

[15] Gu, H. et al. Magnetic cilia carpets with programmable metachronal waves. Nature communications 11, 2637 (2020).

[16] Cui, Z., Wang, Y., Zhang, S., Wang, T. & den Toonder, J. M. Miniaturized metachronal magnetic artificial cilia. Proceedings of the National Academy of Sciences 120, e2304519120 (2023).

[17] Milana, E. et al. Metachronal patterns in artificial cilia for low reynolds number fluid propulsion. Science advances 6, eabd2508 (2020).

[18] Elgeti, J. & Gompper, G. Emergence of metachronal waves in cilia arrays. PNAS 110, 4470–4475 (2013).

[19] Chateau, S., Favier, J., Poncet, S. & d’Ortona, U. Why antiplectic metachronal cilia waves are optimal to transport bronchial mucus. Physical Review E 100, 042405 (2019).

[20] Osterman, N. & Vilfan, A. Finding the ciliary beating pattern with optimal efficiency. Proceedings of the National Academy of Sciences 108, 15727–15732 (2011).

[21] Brumley, D. R., Wan, K. Y., Polin, M. & Goldstein, R. E. Flagellar synchronization through direct hydrodynamic interactions. eLife 3, 1–15 (2014).

[22] Brumley, D. R., Polin, M., Pedley, T. J. & Goldstein, R. E. Metachronal waves in the flagellar beating of Volvox and their hydrodynamic origin. Journal of the Royal Society Interface 12 (2015). URL http://dx.doi.org/10.1098/rsif.2014.1358.

[23] Wan, K. Y. & Goldstein, R. E. Coordinated beating of algal flagella is mediated by basal coupling. PNAS 113, E2784–E2793 (2016).

[24] Quaranta, G., Aubin-Tam, M.-E. & Tam, D. Hydrodynamics versus intracellular coupling in the synchronization of eukaryotic flagella. Physical review letters 115, 238101 (2015).

[25] Salisbury, J., Baron, A., Surek, B. & Melkonian, M. Striated flagellar roots: isolation and partial characterization of a calcium-modulated contractile organelle. The Journal of cell biology 99, 962–970 (1984).

[26] Soh, A. W. et al. Intracellular connections between basal bodies promote the coordinated behavior of motile cilia. Molecular biology of the cell 33, br18 (2022).

[27] Wan, K. Y. Synchrony and symmetry-breaking in active flagellar coordination. Philosophical Transactions of the Royal Society B 375, 20190393 (2020).

[28] Larson, B. T., Garbus, J., Pollack, J. B. & Marshall, W. F. A unicellular walker controlled by a microtubule-based finite-state machine. Current Biology 32, 3745–3757 (2022).

[29] Niedermayer, T., Eckhardt, B. & Lenz, P. Synchronization, phase locking, and metachronal wave formation in ciliary chains. Chaos 18, 37128 (2008). URL 10.1063/1.2956984.

[30] Brumley, D. R., Polin, M., Pedley, T. J. & Goldstein, R. E. Hydrodynamic synchronization and metachronal waves on the surface of the colonial alga Volvox carteri. Physical Review Letters 109, 28–32 (2012).

[31] Pellicciotta, N. et al. Entrainment of mammalian motile cilia in the brain with hydrodynamic forces. PNAS 117 (2020). URL 10.5281/zenodo.3604352.y.

[32] Ringers, C. et al. Novel analytical tools reveal that local synchronization of cilia coincides with tissue-scale metachronal waves in zebrafish multiciliated epithelia. eLife 12, 1–27 (2023).

[33] Guo, H., Man, Y., Wan, K. Y. & Kanso, E. Intracellular coupling modulates biflagellar synchrony. Journal of The Royal Society Interface 18, 20200660 (2021).

[34] Klindt, G. S., Ruloff, C., Wagner, C. & Friedrich, B. M. In-phase and anti-phase flagellar synchronization by waveform compliance and basal coupling. New Journal of Physics 19, 113052 (2017).

[35] Liu, Y., Claydon, R., Polin, M. & Brumley, D. R. Transitions in synchronization states of model cilia through basal-connection coupling. Journal of the Royal Society Interface 15 (2018).

[36] Bruot, N. & Cicuta, P. Realizing the Physics of Motile Cilia Synchronization with Driven Colloids. Annual Review of Condensed Matter Physics 7, 323–348 (2016).

[37] Meng, F., Bennett, R. R., Uchida, N. & Golestanian, R. Conditions for metachronal coordination in arrays of model cilia. PNAS 118 (2021).

[38] Chakrabarti, B., Fürthauer, S. & Shelley, M. J. A multiscale biophysical model gives quantized metachronal waves in a lattice of beating cilia. Proceedings of the National Academy of Sciences of the United States of America 119 (2022).

[39] Hempelmann, F. Zur naturgeschichte von nereis dumerilii aud. et edw. Zoologica (1911).

[40] Özpolat, B. D. et al. The nereid on the rise: Platynereis as a model system. EvoDevo 12, 10 (2021).

[41] Williams, E. A. & Jékely, G. Neuronal cell types in the annelid platynereis dumerilii. Current Opinion in Neurobiology 56, 106–116 (2019).

[42] Fischer, A. H., Henrich, T. & Arendt, D. The normal development of Platynereis dumerilii (Nereididae, Annelida). Front. Zool. 7 (2010).

[43] Jékely, G. et al. Mechanism of phototaxis in marine zooplankton. Nature 456, 395–399 (2008).

[44] Randel, N. et al. Neuronal connectome of a sensory-motor circuit for visual navigation. eLife 3, 1–22 (2014).

[45] Verasztó, C. et al. Ciliomotor circuitry underlying whole-body coordination of ciliary activity in the platynereis larva. eLife 6, 1–25 (2017).

[46] Bezares-Calderón, L. A. et al. Neural circuitry of a polycystin-mediated hydrodynamic startle response for predator avoidance. eLife 7, 1–28 (2018).

[47] Koehl, M. & Hadfield, M. G. Hydrodynamics of larval settlement from a larva’s point of view. Integrative and Comparative Biology 50, 539–551 (2010).

[48] von Dassow, G. & Ellison, C. I. Large-scale ciliary reversal mediates capture of individual algal prey by müller’s larva. Invertebrate Biology 139, e12274 (2020).

[49] Ramirez-San Juan, G.R. et al. Multi-scale spatial heterogeneity enhances particle clearance in airway ciliary arrays. Nature Physics 16, 958–964 (2020).

[50] Purcell, E. M. Life at low Reynolds number. American Journal of Physics 45, 3–11 (1977).

[51] Lauga, E. & Powers, T. R. The hydrodynamics of swimming microorganisms. Reports on Progress in Physics 72 (2009).

[52] Wan, K. Y. et al. Reorganisation of complex ciliary flows around regenerating Stentor coeruleus. Phil. Trans. R. Soc. B 375, 1–9 (2019).

[53] Iatsenko, D. et al. MODA v1.01 (2019). URL 10.5281/zenodo.3470856.

[54] Wan, K. Y., Leptos, K. C. & Goldstein, R. E. Lag, lock, sync, slip: The many ‘phases’ of coupled flagella. Journal of the Royal Society Interface 11 (2014).

[55] Wacker, M. & Witte, H. Time-frequency techniques in biomedical signal analysis. Methods of information in medicine 52, 279–296 (2013).

[56] Wada, Y., Mogami, Y. & Baba, S. A. Modification of ciliary beating in sea urchin larvae induced by neurotransmitters: beat-plane rotation and control of frequency fluctuation. Journal of experimental biology 200, 9–18 (1997).

[57] König, P., Krain, B., Krasteva, G. & Kummer, W. Serotonin increases cilia-driven particle transport via an acetylcholine-independent pathway in the mouse trachea. PloS one 4, e4938 (2009).

[58] Hanasoge, S., Hesketh, P. J. & Alexeev, A. Metachronal motion of artificial magnetic cilia. Soft Matter 14, 3689–3693 (2018).

[59] Kanale, A. V., Ling, F., Guo, H., Fürthauer, S. & Kanso, E. Spontaneous phase coordination and fluid pumping in model ciliary carpets. Proceedings of the National Academy of Sciences 119, e2214413119 (2022).

[60] Mesdjian, O. et al. Longitudinal to transverse metachronal wave transitions in an in vitro model of ciliated bronchial epithelium. Physical Review Letters 129, 038101 (2022).

[61] Von Kenne, A., Bär, M. & Niedermayer, T. Hydrodynamic synchronization of elastic cilia: How surface effects determine the characteristics of metachronal waves. Physical Review E 109, 1–16 (2024).

[62] Solovev, A. & Friedrich, B. M. Synchronization in cilia carpets: Multiple metachronal waves are stable, but one wave dominates. New Journal of Physics 24 (2022).

[63] Chelakkot, R., Hagan, M. F. & Gopinath, A. Synchronized oscillations, traveling waves, and jammed clusters induced by steric interactions in active filament arrays. Soft matter 17, 1091–1104 (2021).

[64] Quillen, A., Peshkov, A., Wright, E. & McGaffigan, S. Metachronal waves in concentrations of swimming turbatrix aceti nematodes and an oscillator chain model for their coordinated motions. Physical Review E 104, 014412 (2021).

[65] Narematsu, N., Quek, R.Chiam, K.-H. & Iwadate, Y. Ciliary metachronal wave propagation on the compliant surface of paramecium cells. Cytoskeleton 72, 633–646 (2015).

[66] Nasi, E. & Tillotson, D. The rate of diffusion of Ca2+ and Ba2+ in a nerve cell body. Biophysical journal 47, 735–738 (1985).

[67] Cortese, D. & Wan, K. Y. Control of helical navigation by three-dimensional flagellar beating. Physical Review Letters 126, 088003 (2021).

[68] Machemer, H. Ciliary activity and the origin of metachrony in paramecium: effects of increased viscosity. Journal of Experimental Biology 57, 239–259 (1972).

[69] Marumo, A., Yamagishi, M. & Yajima, J. Three-dimensional tracking of the ciliate tetrahymena reveals the mechanism of ciliary stroke-driven helical swimming. Communications biology 4, 1209 (2021).

[70] Gheber, L. & Priel, Z. On metachronism in ciliary systems: A model describing the dependence of the metachronal wave properties on the intrinsic ciliary parameters. Cell Motility and the Cytoskeleton 16, 167–181 (1990).

[71] Cheng, Z., Vilfan, A., Wang, Y. & Golestanian, R. Near-field hydrodynamic interactions determine travelling wave directions of collectively beating cilia. Journal of the Royal Society Interface 21 (2024).

[72] Cohen, A. H., Holmes, P. J. & Rand, R. H. The nature of the coupling between segmental oscillators of the lamprey spinal generator for locomotion: A mathematical model. Journal of Mathematical Biology 13, 345–369 (1982).

[73] Boselli, F., Jullien, J., Lauga, E. & Goldstein, R. E. Fluid mechanics of mosaic ciliated tissues. Physical review letters 127, 198102 (2021).

[74] Párducz, B. Ciliary movement and coordination in ciliates. International review of cytology 21, 91–128 (1967).

[75] Feriani, L. et al. Assessing the Collective Dynamics of Motile Cilia in Cultures of Human Airway Cells by Multiscale DDM. Biophysical Journal 113, 109–119 (2017). URL 10.1016/j.bpj.2017.05.028.

[76] Bull, M. S., Kroo, L. A. & Prakash, M. Excitable mechanics embodied in a walking cilium. arXiv preprint 2107.02930 (2021).

[77] Laeverenz-Schlogelhofer, H. & Wan, K. Y. Bioelectric control of locomotor gaits in the walking ciliate euplotes. Current Biology 34, 697–709 (2024).

[78] Hird, C., Jékely, G. & Williams, E. A. Microalgal biofilm induces larval settlement in the model marine worm platynereis dumerilii. Royal Society Open Science 11, 240274 (2024).

