## Supplementary material for "Dynamics and emergence of metachronal waves in the ciliary band of a metazoan larva": SI Text File

### Supplementary Information for:
#### Dynamics and emergence of metachronal waves in the ciliary band of a metazoan larva

Rebecca N. Poon[a,b], Gáspár Jékely[a,c,d], and Kirsty Y. Wan[a,b,*]

[a]Living Systems Institute, University of Exeter, Exeter, EX4 4QD, United Kingdom
[b]Department of Mathematics and Statistics, University of Exeter, Exeter, EX4 4QD, United Kingdom
[c]Biosciences, University of Exeter, Exeter, EX4 4QD, United Kingdom
[d]Centre for Organismal Studies (COS), Heidelberg University, 69120 Heidelberg, Germany

February 14, 2025

#### 1 Automatic cell boundary identification

The image intensity kymographs associated with the *Platynereis* ciliary band metachronal wave appear to be segmented at approximately 30° intervals. These locations coincide with the natural boundaries between adjacent multiciliated cells. The image analysis procedure for automating the identification of these boundaries is summarised in Figure SI 1.

#### 2 Spatial patterning of the metachronal wave

Just like the frequency profile in Figure 1, the wavelength of the metachronal wave is non-uniform over the ciliary band. We can also calculate a wavelength per each cell Figure SI 2a. Most cells contain a maximum of 2 wavelengths, making it impossible to calculate a spatially-resolved average value for the metachronal wavelength, unlike for the beat frequency. Instead, we estimated wavelength from the distance between intensity maxima or minima in the spatial direction, within each cell of the kymograph. Across $n = 21$ larvae, the per-cell wavelength and ciliary beat frequency are negatively correlated, suggesting the wave speed is constant Figure SI 2. For all observed larvae, $f(\theta)$ shows the same 'stepped' profile as in the main text. The binned average shows that there is no anatomical patterning of $f(\theta)$ across different individuals, but all of the larvae show a similar degree of frequency variation. The frequency variation thus does not appear to be anatomically stereotyped Figure SI 3.

#### 3 Immunofluorescence imaging

To confirm that cilia have indeed been ablated and to measure the spacing between ciliary basal bodies, we performed confocal immunofluorescence on larval samples after laser ablation. These larvae were immunostained for acetylated tubulin. Primary antibody: Monoclonal Anti-Tubulin, acetylated antibody produced in mouse (Sigma-Aldrich, Cat# T6793, RRID:AB-477585, 1:250 dilution); and secondary antibody: Goat anti-Mouse Secondary Antibody, Alexa Fluor$^{TM}$ 488, 1:250 dilution). Larvae were imaged on a Zeiss LSM 880 confocal microscope (63x oil immersion objective). The Airyscan superresolution function was used to postprocess the images. Fig. 4(c) shows a $z$-projection of one of the stacks. The ciliary 'stubs' left by the ablation procedure are clearly visible. We measure an average spacing of $(0.57 \pm 0.15)\,\mu$m, and density of $3\,\mu\text{m}^{-2}$. A single cell is $8\,\mu$m tall and $40\,\mu$m wide, so that there are approximately 1000 cilia per cell, or 23,000 cilia in over the 23 cells of the prototroch. The gap between cells in the circumferential direction is $(3.5 \pm 1.0)\,\mu$m, but the gap in the anterior-posterior direction is minimal ($<1\,\mu$m).

#### 4 Creating fire-polished pipettes to hold marine larvae

The tip size of glass micropipettes required for our purposes could be produced directly by the puller, so we outline our protocol here, Fig. 6. We follow the instructions in the Sutter 'Pipette Cookbook' for making holding pipettes, which are generally used for holding cells or blastocysts stationary for microinjections.

The puller is programmed to generate a very long taper, which is then 'snapped off' at the correct location to give the desired tip diameter. (The precise values used for our puller were: Heat = Ramp + 10; Pull = 85; Velocity = 90; Time = 90; Pressure = 200. Our puller is fitted with a $3 \times 3$mm box filament.) For our purposes, the ideal inner diameter after this step is $\approx$60 µm. A clean break can be achieved by scoring a mark across the taper at the correct height using the rough face of a small ceramic tile (from Sutter Instruments), and then pushing on the tip above the score mark so that the pipette breaks cleanly at the score point at a 90° angle. It is difficult to score and break off the long taper at a consistent location, so the initial tip diameter generated by this step is quite variable. Additionally, the break is not always clean and at 90°. It is usually possible to make a second break on the same pipette, but this will naturally produce a larger tip diameter.

This broken end produced by this first step is very sharp, so the edges must be rounded off by a process of repeated, even heating known as fire polishing. If this step is omitted, the sharp walls can puncture the soft walls of the larval cells. Fire polishing involves holding the very tip of the broken-off pipette stationary inside the filament of the puller and heating it for short periods of time until the edges are sufficiently round.

The degree of melting achieved by each heating event must be controlled by changing the heat and time settings on the puller. (For fire polishing, we use Ramp$\leq$Heat$\leq$Ramp+40; Pull = 0; Velocity = 0; 150$\leq$Time$\leq$200; Pressure = 500.) As the walls at the pipette tip 'round out' due to the heating, the inner diameter of the pipette also decreases. By performing multiple heating cycles, an initial inner diameter of as high as 90 µm can be melted down to an inner diameter of 30 µm, although this produces a tip with very thick walls. Additionally, if the tip is heated too much, the tip closes and the pipette becomes unuseable.

The ideal inner diameter for holding *Platynereis* larvae was found to be 20−30 µm and the ideal wall thickness is 30−40 µm. If the inner diameter is too small, it is difficult to apply sufficient force to hold the larva stationary. If the inner diameter is too large, the suction pressure is applied on a very large area and causes unnecessary levels of cell deformation.

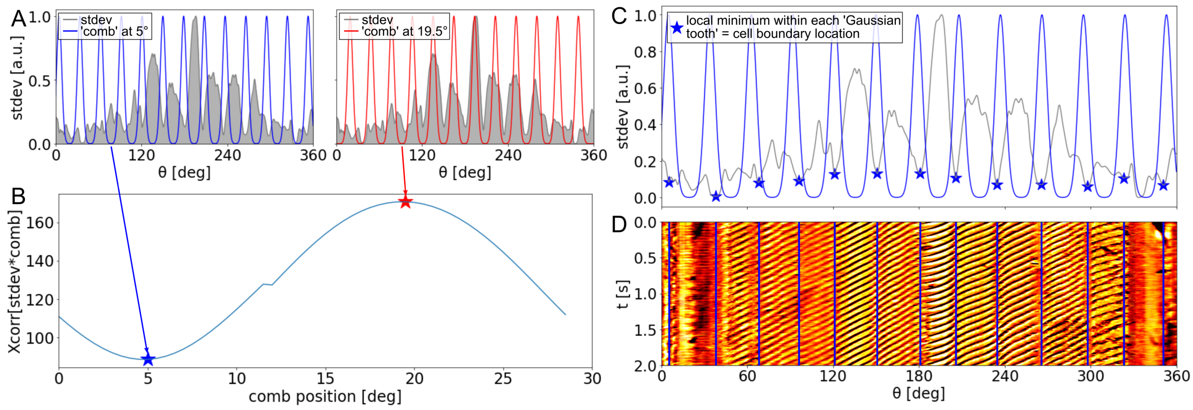


Fig. SI 1: **Method for finding the locations of cell boundaries from the discontinuities in the wave intensity kymograph**. Gap locations are constrained to be spaced at $\approx 30°$. (A) The standard deviation of the image intensity over time at each location around the ciliary band has a minimum value at each cell boundary, due to the absence of cilia producing a periodically varying signal at that location. We create a 'comb' of multiple Gaussians, with the peaks spaced at approximately 30°. We shift the comb to different locations around the band, and calculate (B) the cross correlation of the standard deviation with the combs at different locations. The minimum (and maximum) values of this cross-correlation indicate where the comb is 'best positioned' over the minima (and maxima) of the standard deviation function. (C) We use the comb positioned at the 'minimum value location'. Within each individual 'Gaussian tooth' of the comb, we find the location of the local minimum in the standard deviation, and designate these as the cell boundaries. (D) visual inspection of the kymograph confirms accurate location of the boundaries. The 'teeth' of the comb were generally spaced at $(30 \pm 2)°$ as needed to correctly locate the boundaries, due to variations in individual cell structure and imaging orientation. The Gaussian teeth can be widened to 'weaken' the restriction of approximately equal cell spacing.

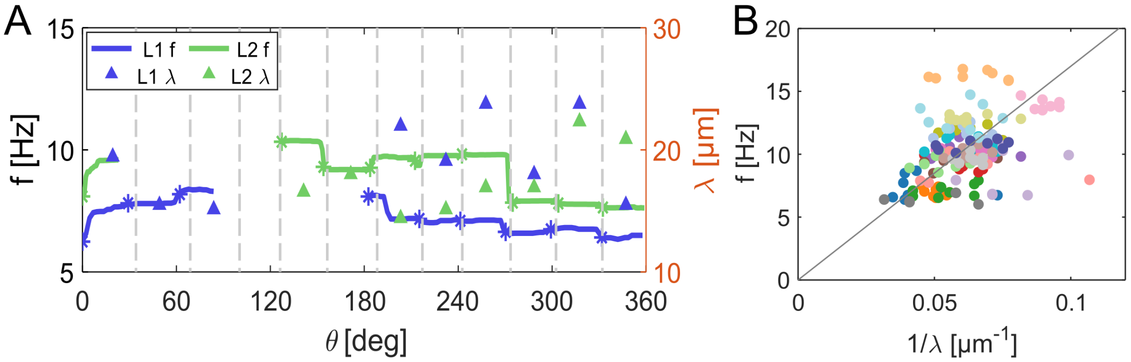


Fig. SI 2: **Wavelength profile along the ciliary band and wavelength-frequency relationship.** (A) Frequency and wavelength $\lambda$ for two larvae (further to that shown in Fig. 1(I)), showing inter-cellular variation in the wavelength. (B) Frequency against $\lambda^{-1}$ for all of the individual cells of the dataset, colour coded by larva number. A group of points of the same colour shows all of the cells for a single individual, and shows no correlation. However, across all of the individuals of the dataset there is a weak positive correlation (Pearson correlation coefficient = 0.4), indicating that the average wave speed is similar for multiple individuals. Taking the average value all points gives a wave speed of $170\,\mu\mathrm{m\,s^{-1}}$: the grey line shows where $f\lambda = 170\,\mu\mathrm{m}$.

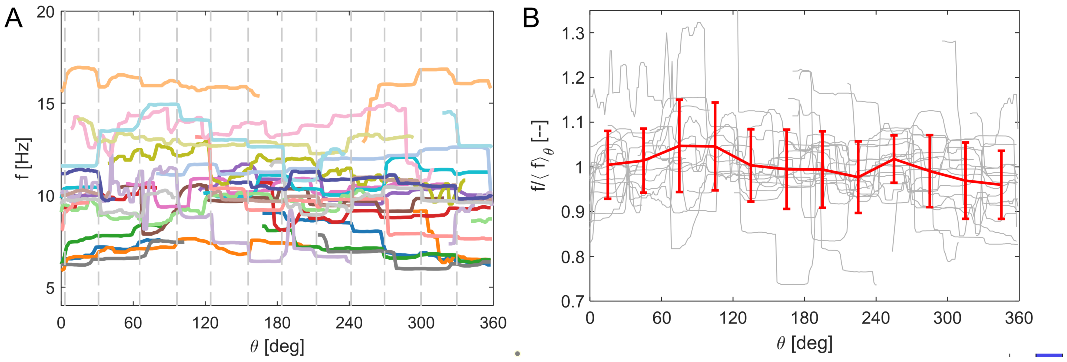


Fig. SI 3: **Spatially-resolved frequency measurements across multiple larvae.** (A) The $f(\theta)$ profile for 21 individuals. Data is omitted within the pipette, and $\theta$ is normalised to the dorsal point of each individual. All individuals show the same 'stepped' profile and the average larval beat frequency is $(10 \pm 2)$ Hz. (B) We investigated the possibility of this variation being a stereotyped function of the position around the larva by plotting the frequencies (normalised by the average frequency for each individual) (grey), and the cell binned average across all 21 individuals (red). Error bars show standard deviation within the bin.

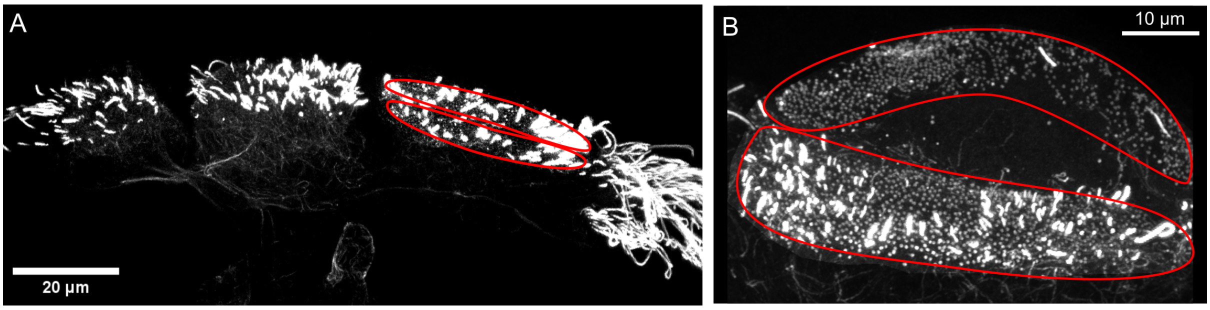


Fig. SI 4: **Confocal immunofluorescence imaging of larval ciliary band cells that have been partially de-ciliated by laser ablation**. The red lines show the approximate locations of an anterior-posterior pair of cells on each image. (A) shows four intact cell pairs from the ventral side of a laser-ablated larva. The three leftmost cells have been mostly de-ciliated, whilst the rightmost cell retains its long cilia. The larva has been stained for acetylated tubulin. We use this image to measure the size of the gap between cilia across a cell boundary. (B) shows a higher magnification view of an anterior-posterior pair of cells that has been almost entirely de-ciliated, leaving the ciliary basal bodies visible as dots. The pair of cells has split apart during sample preparation, leaving a wide gap between them. We use this image to measure the spacing and density of the cilia within a single cell.

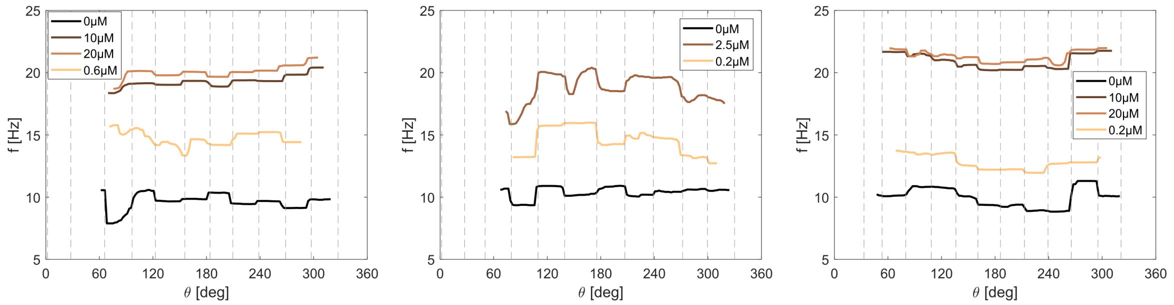


Fig. SI 5: **Wash-out of extracellular serotonin.** After increasing the serotonin concentration, the serotonin can be 'washed out' by repeatedly replacing known volumes of the surrounding solution with fresh seawater (0 µM serotonin). Plotting $f(\theta)$ for three individuals for increasing serotonin concentration points as well as the subsequent 'washed out' profile shows that the frequency recovers a lower value when the concentration decreases. The fact that the 'washed out' frequency is still higher than the original frequency may be due to the impossibility of reducing the extracellular concentration to 0 µM simply by replacing fluid; or a residual higher concentration of intracellular compared to extracellular serotonin; or some larval response to the process of replacing the surrounding fluid.

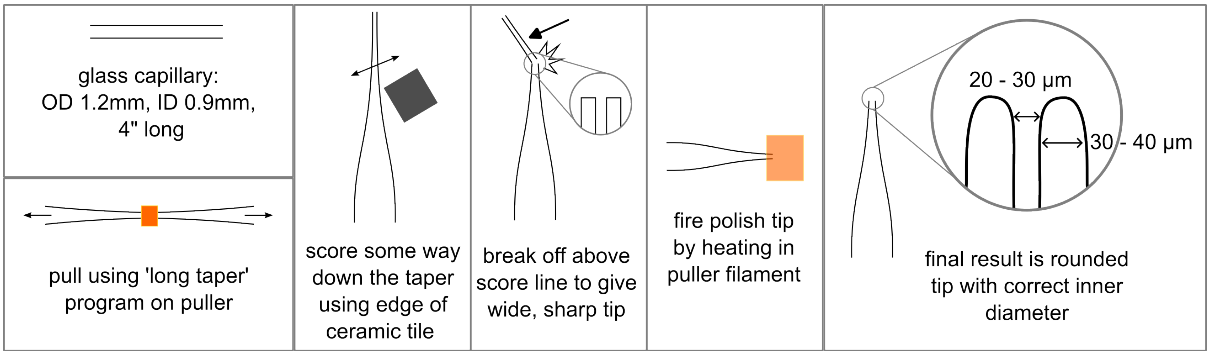


Fig. SI 6: **Micropipette fabrication and fire-polishing procedure.** Process for making a micropipette suitable for holding larvae of ≈ 100–500 µm size. We use a digital pipette puller to pull a 'long taper' pipette. We then score and break off the long taper to generate a large tip with sharp edges. We fire polish the tip to round off the edges – this process can also be used to reduce the inner diameter. This process generates a rounded pipette tip with our desired inner diameter.

#### List of SI Video captions

- Movie S1: Visualisation of the ciliary band metachronal wave in the prototroch on a pipette-held *Platynereis dumerilii* larva ($24 - 36$ hpf) .
- Movie S2: A small gap in the ciliary band was created by laser ablation. The video shows the dynamics of the metachronal wave before, during and after the ablation process.
- Movie S3: Progressive laser ablations at different locations along the ciliary band of the same larva (time point labels refer to Fig. 4A). The video shows a region of around 3-cell widths, as more and more cilia are removed. In the last time point, only one small patch of cilia from one cell remains in the whole ciliary band. A metachronal wave continues to propagate within this one cell.
- Movie S4: Closures are spontaneous whole-body events and transient ciliary arrest associated with calcium signalling. Here, we focus on a single closure event where the entire ciliary band is visible. The closure begins with near-simultaneous arrest of all cilia, a quiescent period where all cilia are arrested, and finally sequential (but fast) recovery of the metachronal wave, where local spatial patterning is observed immediately as the cilia restart their beating.
